# Genome-wide Analysis Reveals High Genomic Diversity and Panmixia in Bay Pipefish (*Syngnathus leptorhynchus*) from Coos Bay, Oregon

**DOI:** 10.64898/2025.12.31.696303

**Authors:** Mark C. Currey, Micah A. Woods, Vithika Goyal, Susan Bassham, Hope M. Healey, William A. Cresko

**Author notes:** Department of History and Philosophy of Science, University of Cambridge, Cambridge, United Kingdom (Micah A. Woods). Email: (M.A.W.); (V.G.); (S.B.); (H.M.H.). Send correspondence to this address (M.C.C.). Send correspondence to this address (W.A.C.).

## Abstract

The over 300 species of fishes in the charismatic family Syngnathidae (seahorses, pipefish, pipehorses, and seadragons) exhibit remarkable morphological and reproductive adaptations. Most syngnathid species inhabit shallow coastal regions where they are vulnerable to human activities, including habitat degradation and collection for use in traditional medicine and the aquarium trade. A gap remains in our understanding of within-species genetic diversity and population structure for most of the species in this family—information that is crucial for establishing baselines to detect future human-driven changes. Here, we investigate fine-scale population structure and genomic diversity in the bay pipefish (*Syngnathus leptorhynchus*), one of the relatively few temperate lineages within the Syngnathidae. Bay pipefish are habitat specialists that are closely associated with eelgrass (*Zostera marina*) along the Pacific coast of North America. Due to the patchy and changing nature of eelgrass habitat, bay pipefish provide an instructive case for using population genomics to explore patterns of connectivity in a dynamic coastal environment. We sampled 116 individuals from two locations separated by approximately 17 km in the Coos Bay estuary (Oregon, USA) and used restriction site-associated DNA sequencing (RAD-seq) to discover high and consistent genetic diversity at both sampling locations, with no indication of inbreeding. Genome-wide patterns of genetic variation showed minimal differentiation between the sampling sites, with levels of divergence indistinguishable from those expected under random sampling. Linkage disequilibrium was low and declined rapidly, and estimates of effective population size indicate a large, well-connected population within Coos Bay. These results establish a crucial genomic baseline for understanding how bay pipefish populations may respond to ongoing changes in their eelgrass habitat and offer a genomic framework for conservation and comparative studies across the Syngnathidae.

## INTRODUCTION

Understanding how habitat specialists maintain genetic connectivity across fragmented or dynamic habitats is critical for predicting their responses to environmental change, particularly in marine systems where connectivity patterns remain poorly understood for many taxa (Cowen and Sponaugle, 2009; Lowe and Allendorf, 2010). The family Syngnathidae, which includes over 300 species of pipefish, seahorses, pipehorses, and seadragons, is characterized by specialized morphology and strong habitat associations (Dawson, 1985; Mobley et al., 2011). Most syngnathid species are adapted to shallow coastal and estuarine environments that are increasingly impacted by outcomes from human activities, including habitat degradation, pollution, and climate-driven shifts in temperature and salinity (Foster and Vincent, 2004). Many of these species also face direct human pressures through harvesting for traditional medicine, the aquarium trade, and scientific research (Vincent et al., 2011).

Despite their ecological, scientific, and cultural importance, within-species patterns of genetic diversity, connectivity, and population structure remain poorly characterized for much of the Syngnathidae (Mobley et al., 2011; Vincent et al., 2011). These knowledge gaps limit our ability to assess population resilience to environmental change and exploitation. Establishing robust baseline measurements of genomic diversity and spatial genetic structure is therefore critical for detecting future shifts in population status and for informing effective conservation and management strategies for syngnathid and other vulnerable fish taxa. Furthermore, the small number of population genetic studies in syngnathids have revealed variable findings. Specialized habitat dependence and limited dispersal have been linked to fine-scale genetic structure in many syngnathids (Teske et al., 2003; Lourie et al., 2005; Stiller et al., 2017; Bertola et al., 2020), though patterns vary considerably. Some species show strong population structure at small spatial scales (Partridge et al., 2012), while others exhibit surprisingly broad genetic connectivity despite being distributed over large geographic distances (Mobley et al., 2010; Braga Goncalves et al., 2017; Woodall et al., 2018). These contrasting patterns suggest that connectivity in habitat-specialist syngnathids depends on complex interactions between life history traits, habitat configuration, and species-specific dispersal behavior.

The bay pipefish (*Syngnathus leptorhynchus*) provides an exemplary syngnathid system for examining population genetic patterns. Despite having one of the broadest distributions among *Syngnathus* species—from Bahía Santa María, Baja California to Prince William Sound, Alaska (Fritzsche, 1980; Orsi et al., 1991)—bay pipefish are strongly associated with eelgrass (*Zostera marina*) throughout their life cycle (Adams, 1976; Fritzsche, 1980). Previous studies of bay pipefish have examined population structure at different spatial scales, with findings that are complementary but leave some uncertainty about the factors governing gene flow. Broad-scale phylogeographic work along the west coast of North America, spanning hundreds to thousands of kilometers, documented clear genetic breaks among distant coastal regions (Louie, 2003; Wilson, 2006) whereas fine-scale analyses in British Columbia’s Barkley Sound detected measurable divergence (F_ST_ = 0 - 0.017) among populations separated by a few to tens of kilometers, with seascape complexity restricting gene flow and creating distinct genetic clusters (de Graaf, 2006). These studies leave open the question of whether fine-scale structure could develop in topographically simple, continuous estuaries where physical barriers to dispersal are largely absent. As eelgrass habitat specialists with limited swimming ability, bay pipefish could exhibit restricted dispersal even in continuous habitats, yet the dynamic and patchy nature of eelgrass meadows could alternatively promote high connectivity through repeated colonization and mixing.

Previous bay pipefish studies relied on mitochondrial or microsatellite markers, limiting inferences about genome-wide diversity and genetic connectivity. Here, we use restriction site-associated DNA sequencing (RAD-seq), enabled by the recent assembly of a reference genome, to examine fine-scale population structure and genetic diversity genome-wide in bay pipefish within the Coos Bay estuary, Oregon—a continuous estuarine system where topographic barriers are minimal. We sampled individuals from two locations approximately 17 km apart and characterized genetic variation in thousands of single nucleotide polymorphisms (SNPs) to assess patterns of population structure, genetic diversity, kinship, and effective population size. We assessed whether bay pipefish exhibit (1) fine-scale genetic structure reflecting limited dispersal and habitat specialization, as observed in more topographically complex regions, or (2) panmixia reflecting the continuous nature of an estuarine environment that has potentially high connectivity within eelgrass meadows.

This study establishes a baseline understanding of the genetic diversity and structure of bay pipefish in Coos Bay and offers a point of reference for future monitoring and conservation efforts, as well as a useful comparison for other estuarine systems along the west coast. More broadly, this work contributes to a growing understanding of how spatial scale, habitat continuity, and life history traits shape genetic structure in syngnathid fishes and informs future conservation and sampling strategies in estuarine environments.

## MATERIALS AND METHODS

### Pipefish collections

Bay pipefish were collected by beach seine in the summer of 2023 from two sites in the Coos Bay estuary: “Causeway” and Valino Island (Fig. 1 and Table 1). At each site, beach seines were pulled through areas containing eelgrass beds. We collected 50 adult fish from the Causeway location and 66 from Valino Island. Fish were placed in cooled, aerated seawater for transport until euthanized with tricaine methane sulfonate (MS-222) and immediately preserved in 95% ethanol. GPS coordinates of sampling locations were obtained from Google Earth (Table 1). All research involving vertebrate animals was approved by the University of Oregon Institutional Animal Care and Use Committees under protocol number AUP-20-23. Fish were collected under Oregon Department of Fish and Wildlife scientific taking permit number 26987.

**Figure 1.**
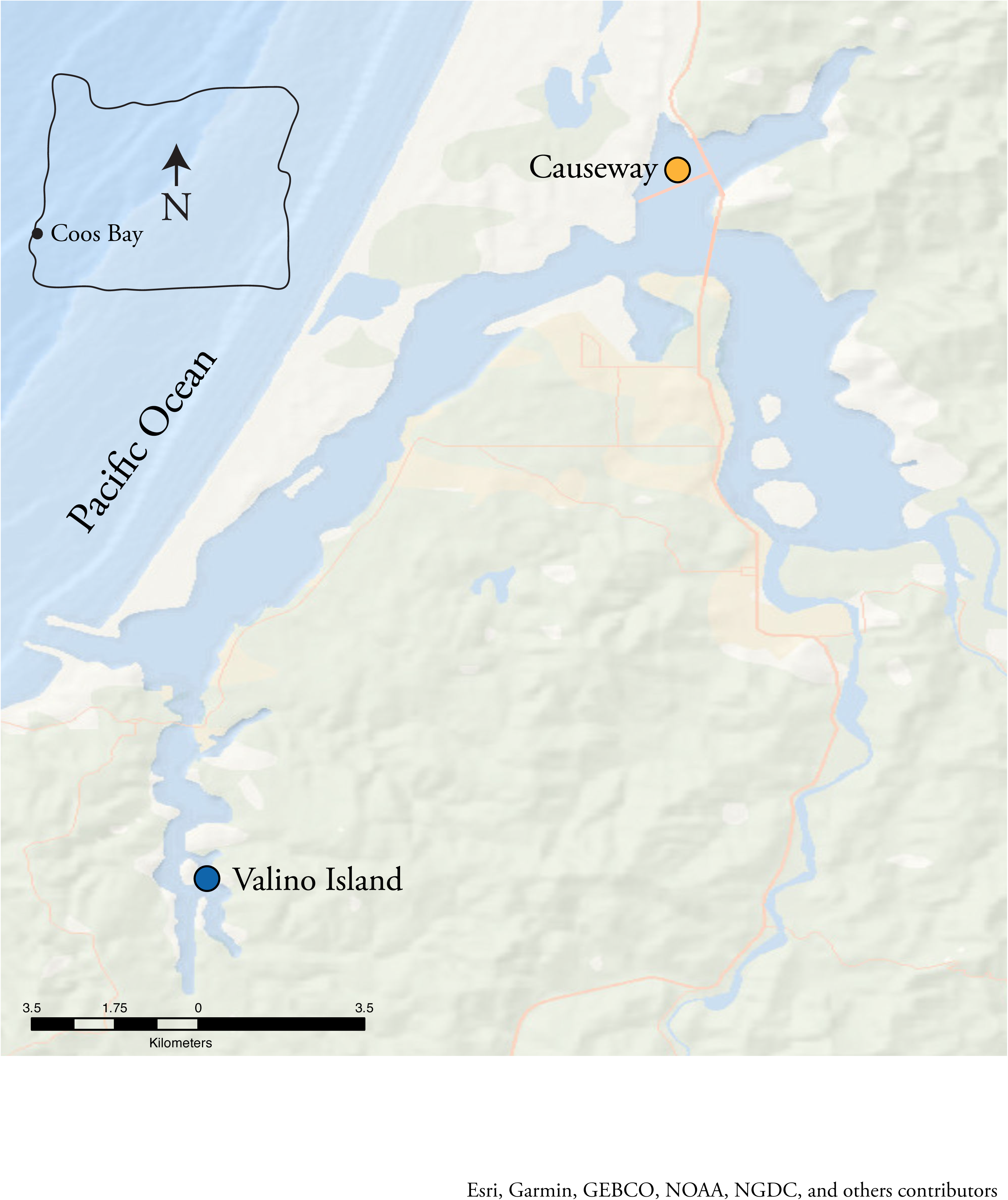
Sampling locations for bay pipefish in Coos Bay, Oregon. The blue point indicates Valino Island, while the orange point marks the Causeway site, positioned adjacent to the Trans Pacific Pkwy.

**Table 1.** Sampling site information. This table presents collection details for two sampling sites. Columns show sample site name, GPS coordinates, elevation in meters and feet, distance to ocean in kilometers and miles, collection dates, and total number of samples collected at each site.

| Sample Site | Sample site GPS coordinates | Elevation (m/ft) | Distance to ocean (km/miles) | Collection dates | Total samples |
| --- | --- | --- | --- | --- | --- |
| Causeway | 43°26'46"N<br>124°13'23"W | 0.52/2 | 16.24/10.09 | September 14, 2023 | 50 |
| Valino Island | 43°18'45"N<br>124°18'55"W | 1.96/6 | 7.64/4.75 | September 14, 2023 | 66 |

### DNA extraction and quality control

Genomic DNA was purified from fin and other body tissues using a column-based DNeasy Blood & Tissue Kit (Qiagen) or by tissue digestion in Qiagen ATL buffer with proteinase K followed by DNA isolation using Omega Tech paramagnetic beads which were employed for high-throughput processing of samples. To avoid potential sampling bias, samples from each location were processed using both extraction methods. All samples underwent an RNaseA treatment. Purified DNA was quantified using a Qubit fluorometer and dsDNA Broad Range Assay Kit. A subset of samples was assessed for DNA purity using a NanoDrop spectrophotometer, and for integrity *via* agarose gel electrophoresis. Both assessments indicated high-quality DNA with minimal contamination. DNA was then normalized in concentration to ensure equal representation of each individual in the RAD-seq library.

### RAD library construction and genotyping

RAD-seq libraries were constructed for 116 fish using the restriction endonuclease *SbfI*-HF (NEB), following established protocols (Catchen et al., 2013a). Sequencing was performed on an Illumina NovaSeq 6000 platform with a sp100 cycle, generating 135-nucleotide reads, including inline 7 nt barcodes. Raw reads were de-multiplexed and filtered by quality score using default settings in the process_radtags module from the Stacks v2.68 pipeline (Catchen et al., 2013b). Briefly, the reads were first checked for intact barcodes and RAD cut sites and then demultiplexed. Next, a sliding window approach (default window size of 15% of read length) was used to assess average quality scores, discarding reads where the score dropped below a Phred score of 10 (corresponding to 90% probability of base call accuracy). Sequencing and quality filtering yielded approximately 500 million initial reads with a little over 373 million sequences retained after passing quality filters (Supplemental Table 1). These high-quality reads were then aligned to the *Syngnathus leptorhynchus* genome (GenBank assembly accession: GCA_XXXXXXXXX, to be assigned upon release) using GSNAP version 2024-05-07 (Wu and Watanabe, 2005), allowing for up to five nucleotide mismatches and gap lengths of two nucleotides. Only reads with unique alignments were retained.

Reads were then processed through the Stacks pipeline (modules: ref_map.pl and populations) to construct a catalog and to call genotypes. Individual samples were filtered using VCFtools (Danecek et al., 2011) based on the following conservative, quality-focused criteria. To ensure that only high-confidence genotypes were retained, samples could not have a missing data rate >12%, exhibit excess heterozygosity or high inbreeding (F < –0.10 or F > 0.25), or show a combination of >10% missing data and moderate heterozygosity deviations (F < –0.05 or F > 0.20). This combined filtering removed 36 individuals in total—14 from the Causeway site and 22 from the Valino Island site—leaving 36 individuals from Causeway and 44 from Valino Island for downstream analyses.

A whitelist (i.e., a list of locus IDs to retain in downstream analyses) of loci including reads from only the first 22, which are likely the chromosomes in this species, was then used in a second populations run. Separate sampling locations were retained in the data structure to enable population structure analyses. For their inclusion, we required that loci be present in at least 80% of individuals within each population (-r 0.8), they be present in at least 80% of all individuals across the study (-R 0.8), have a minimum minor allele frequency of 0.05 (--min-maf 0.05), and, using the --write-single-snp flag, only one SNP per RAD read was retained. This resulted in 6,074 variant sites (Supplemental Table 2). A third populations run was conducted using individuals from both sampling locations combined. The same filtering criteria were applied, except the -r flag was omitted because we collapsed the populations into one. This resulted in 6,310 variant sites (Supplemental Table 2). These two whitelisted sets represent average genome-wide marker densities of ∼20 and ∼21 markers per megabase, respectively.

### Population genetic analysis of diversity and structure

The populations program within the Stacks framework was used to calculate per-locus and genome-wide averages of nucleotide diversity (π), heterozygosity, F_IS_, and F_ST_.

To assess the significance of genome-wide genetic differentiation, we applied two complementary statistical approaches that test distinct null hypotheses. First, permutation testing evaluated whether the observed genome-wide F_ST_ exceeded expectations under panmixia (no population structure). Individuals were randomly reassigned to the two sampling locations while preserving the original sample sizes, and genome-wide F_ST_ was recalculated for each permutation. This procedure was repeated 100 times to generate a null distribution of F_ST_ values. Statistical significance was calculated using the formula *p* = (r + 1) / (n + 1), where *r* is the number of permuted F_ST_ values greater than or equal to the observed value, and *n* is the number of permutations.

Second, bootstrap resampling assessed whether the observed genome-wide F_ST_ could be explained by sampling variance among loci. From the quality-filtered dataset of 6,074 SNPs, we generated 1,000 bootstrap replicates by randomly sampling 1,000 loci with replacement. For each replicate, genome-wide F_ST_ was recalculated using the same parameters as the original analysis in *Stacks*. The resulting distribution was summarized by its mean, median, standard deviation, and 95% confidence interval. The percentile rank of the observed F_ST_ within this distribution was used to assess whether the observed value fell outside the range expected from sampling variance alone.

Output files from the Stacks populations program were generated under two grouping scenarios. First, we analyzed the two sampling sites separately to test for population structure, calculate pairwise divergence (F_ST_), and assess kinship patterns. Second, because preliminary analyses indicated panmixia (see Results), we combined both sites into a single population for analyses that assume random mating: genome-wide linkage disequilibrium (LD) patterns and effective population size (Ne) estimation.

To visualize and investigate the major axes of genetic variation, a principal component analysis (PCA) was performed using Genodive v3.04 software (Meirmans, 2020) employing a covariance method on individual genotype data from the two collecting locations (Causeway and Valino Island). The analysis utilized the populations.structure file generated by Stacks (Catchen et al., 2013b), which contained all 6,074 markers, to calculate genetic principal components to assess population differentiation. Principal component plots were generated in R v4.4.0 (R Core Team, 2024) using *ggplot2* v3.5.2 (Wickham, 2016). The first two components were visualized with their variance explained, and a density plot of PCA scores was also produced using *ggplot2*.

Population genetic structure was investigated using STRUCTURE v2.3.4 (Pritchard et al., 2000), and the same populations.structure file as in the PCA analysis, to assess the number of distinct genetic clusters among individuals from the two collecting locations. For each value of *K* (number of genetic groupings), five independent runs were performed using 100,000 burn-in steps followed by 100,000 Markov Chain Monte Carlo replicates. The range of *K* values tested was 1 to 2 to visualize genetic clustering. Results from multiple runs were processed and visualized using CLUMPAK (Kopelman et al., 2015) to assign individuals to genetic clusters.

Genome-wide patterns of differentiation were assessed using pairwise *F_ST_* estimates calculated by the populations module of Stacks for all retained SNPs between the two collecting locations. To visualize genomic patterns of differentiation, *F_ST_* values were plotted against genomic position using both individual SNP point estimates and smoothed trend lines to identify genomic regions of elevated or reduced differentiation. All visualizations were created using R.

### Kinship Estimation

To assess relatedness among individuals, pairwise kinship coefficients were estimated using the Method of Moments (MoM) in the R package SNPRelate v1.38.1 (Zheng et al., 2012). Prior to analysis, VCFs from Stacks were filtered to retain only biallelic SNPs with ≤25% missing data. Linkage disequilibrium (LD) pruning was applied using a 10 kb sliding window and an LD threshold of *r²* < 0.2 to ensure marker independence. Kinship was calculated for all individual pairs (630 within Causeway, 946 within Valino Island, and 1,584 between locations; total = 3,160 pairs). Results were formatted as symmetric matrices and pairwise tables for downstream analysis. Kinship values were interpreted using standard thresholds (e.g., ≥0.45: identical; ≥0.22: parent-offspring/full siblings; ≥0.11: half-siblings/grandparent-grandchild; ≥0.055: first cousins; ≥0.027: second cousins; >0: distant relatives; ≤0: unrelated). Heatmaps and summary plots were used to visualize relatedness patterns within and between sites using the pheatmap v1.0.13 package in R (Kolde, 2025).

### Genome-wide Linkage Disequilibrium and LD Decay Analysis

Genome-wide linkage disequilibrium (LD) was calculated using PLINK2.0 (Chang et al., 2015) with the flags --r2-phased, cols=+dprimeabs, --inter-chr, and --ld-window-r2 0 to generate both r² and D’ values for all SNP pairs. To visualize LD, values were aggregated into 1 Mb genomic bins using maximum aggregation for r² and mean aggregation for D’. Symmetric heatmaps displaying 287 bins (covering 91% of the genome) across 22 chromosomes were generated using ggplot2.

LD decay was examined using two complementary approaches: a fine-scale analysis (≤500 kb with 10 kb bins) to capture local recombination patterns, and a broad-scale analysis (≤5 Mb with 100 kb bins) to reveal longer range trends. For each distance bin, mean LD values were calculated, excluding bins containing fewer than five SNP pairs and plotted along with raw pairwise LD values (as transparent points) and binned means (as solid lines and points), illustrating the underlying data distribution alongside smoothed trends. Plots were created with R and ggplot2.

The correlation between r² and D’ was calculated across all intra-chromosomal SNP pairs within 5 Mb using Pearson correlation. Background LD levels were defined as the mean r² and D’ values for SNP pairs separated by >1 Mb, representing the baseline level of residual linkage disequilibrium. Decay to background levels was determined using fine-scale 2 kb bins up to 200 kb distance, identifying the first distance bin where LD values dropped to within 10% of the background threshold. Distance-controlled relationships were assessed using linear regression models: (1) r² ∼ D’, (2) r² ∼ D’ + log(distance), and (3) r² ∼ D’ × log(distance), with model improvements evaluated using ANOVA. Partial correlation between r² and D’ was calculated while controlling for log-transformed physical distance using the ppcor v1.1 (Kim, 2015) package in R.

Potential regions of elevated LD on chromosome 1 (Chr 1) were identified using sliding 1 Mb windows. For each window containing ≥10 SNP pairs, mean r² and D’ values were calculated and compared to genome-wide thresholds defined as the mean plus two standard deviations of values from all other chromosome groups. Windows exceeding either threshold were considered elevated, and contiguous elevated windows defined region boundaries. LD magnitude comparisons between the identified elevated region and genome-wide averages (excluding all Chr 1 data) were expressed as fold-changes and absolute differences to quantify the degree of elevation.

All analyses were restricted to intra-chromosomal SNP pairs to avoid confounding effects from inter-chromosomal associations, and statistical significance was assessed at α = 0.05. Analyses were conducted using data.table v1.17.8, dplyr v2.5.1, and ggplot2 v3.5.2 (Wickham et al., 2023; Barrett et al., 2025).

### Contemporary Effective Population size (Ne)

Contemporary effective population size (Ne) was estimated using two complementary approaches to provide cross-validation and increase confidence in the results. These methods capture different aspects of population genetic variation: the allele frequency variance method estimates Ne from the distribution of allele frequencies across loci, while the linkage disequilibrium (LD) method relies on non-random associations between alleles at different loci. Because the two methods are based on independent principles and assumptions—one focusing on allele frequency variance and the other on inter-locus associations—they provide robust and complementary insights into contemporary Ne.

To ensure independence among markers, linkage disequilibrium (LD) pruning was performed using PLINK v2.0 with the --indep-pairwise 50 10 0.2 algorithm, which examines SNPs in sliding windows of 50 markers, shifts by 10 SNPs per iteration, and removes one SNP from each pair exhibiting LD with r² > 0.2. This retained 5,676 independent SNPs from the original dataset for all Ne analyses.

Ne was first estimated using the allele frequency variance method described by Nei and Tajima (Nei and Tajima, 1981), which calculates effective population size based on the variance in allele frequencies across loci. Specifically, Ne is estimated using the formula Ne = *n* / (2 × Var(*p*)), where *n* is the sample size, and Var(*p*) is the variance in minor allele frequencies. To quantify uncertainty, 1,000 bootstrap replicates were performed by resampling loci with replacement from the pruned SNP dataset, and 95% and 90% confidence intervals were calculated from the resulting distribution of Ne values. Analysis was implemented in R using custom scripts with the data.table package (Barrett et al., 2025).

A second estimate of Ne was obtained using the linkage disequilibrium method implemented in NeEstimator v2 (Do et al., 2014), which infers effective population size based on correlations between alleles at different loci. To evaluate sensitivity to rare alleles, Ne was estimated using multiple critical allele frequency thresholds (Pcrit = 0.05, 0.02, and 0.01), with Pcrit = 0.02 selected as the primary estimate following standard recommendations for balanced precision and bias. Genotype data were converted from PLINK to GenePop format using custom R scripts, and parametric 95% confidence intervals were calculated from the standard errors reported by NeEstimator.

## RESULTS

### Notable genetic variation within and among collecting locations

Of the 6,074 variant sites identified in Stacks, none carried alleles unique to the Causeway collection. Only one variant site had an allele private to the Valino Island collection, which likely reflects sampling variation rather than a fixed presence/absence polymorphism at that locus. Genome-wide averages for the Causeway site were as follows: expected heterozygosity (H_e_) 0.293, observed heterozygosity (H_o_) 0.257, nucleotide diversity (π) 0.298, and inbreeding coefficient (F_IS_) 0.130. For the Valino Island site, the corresponding values were H_e_ 0.297, H_o_ 0.259, π 0.301, and F_IS_ 0.136. When both locations were analyzed together, genome-wide averages were H_e_ 0.300, H_o_ 0.256, π 0.302, and F_IS_ 0.149 (Supplemental Table 2).

### Little to no observed population structure between collecting locations

PCA of 80 individuals revealed that the first two principal components captured 2.0% and 1.6% of the total genetic variance (Fig. 2, Supplemental Figure 1). Individuals from the two sampling locations largely overlapped in PCA space, with no clear clustering by site, indicating minimal genetic differentiation across the 17 km separating the habitats. STRUCTURE analysis revealed a similar pattern, with each individual showing nearly equal assignment probabilities to two hypothesized populations (K = 2), regardless of their collection site (Fig. 3). Despite using thousands of genomic markers—which provide substantially greater resolution than previous analyses using relatively fewer microsatellite markers—no spatial genetic structure was detected. The lack of clear assignment of individuals to distinct genetic clusters further supports the conclusion that the individuals collected from the two locations represent a single, well-mixed population rather than genetically differentiated groups.

**Figure 2.**
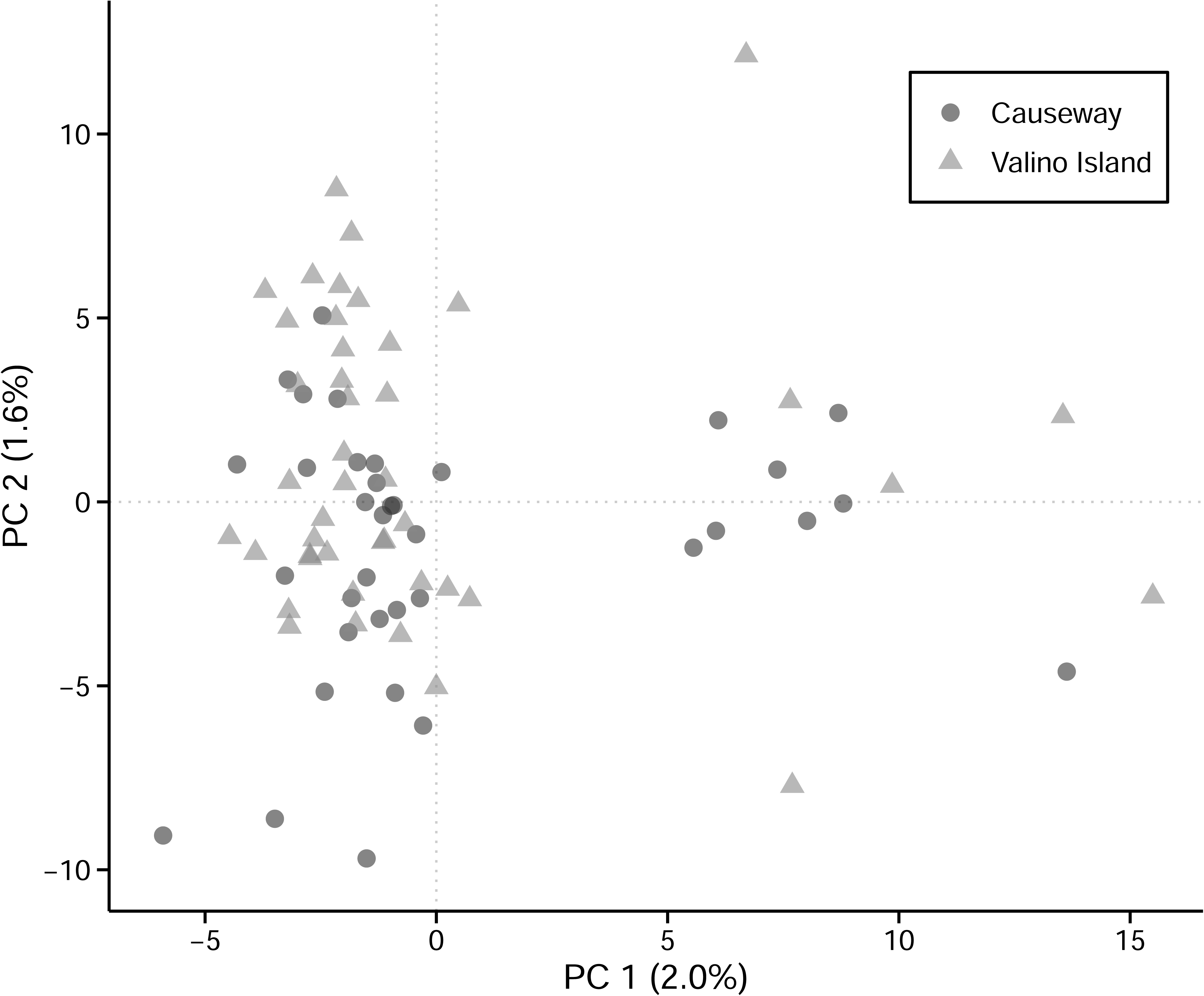
Principal Component Analysis of genetic variation between two populations. PCA plot showing genetic differentiation between Causeway (dark gray circles) and Valino Island (light gray triangles) individuals. PC1 (2.0%) and PC2 (1.6%) explain 3.6% of the total genetic variance. Each point represents an individual, with color and shape distinguishing populations.

**Figure 3.**
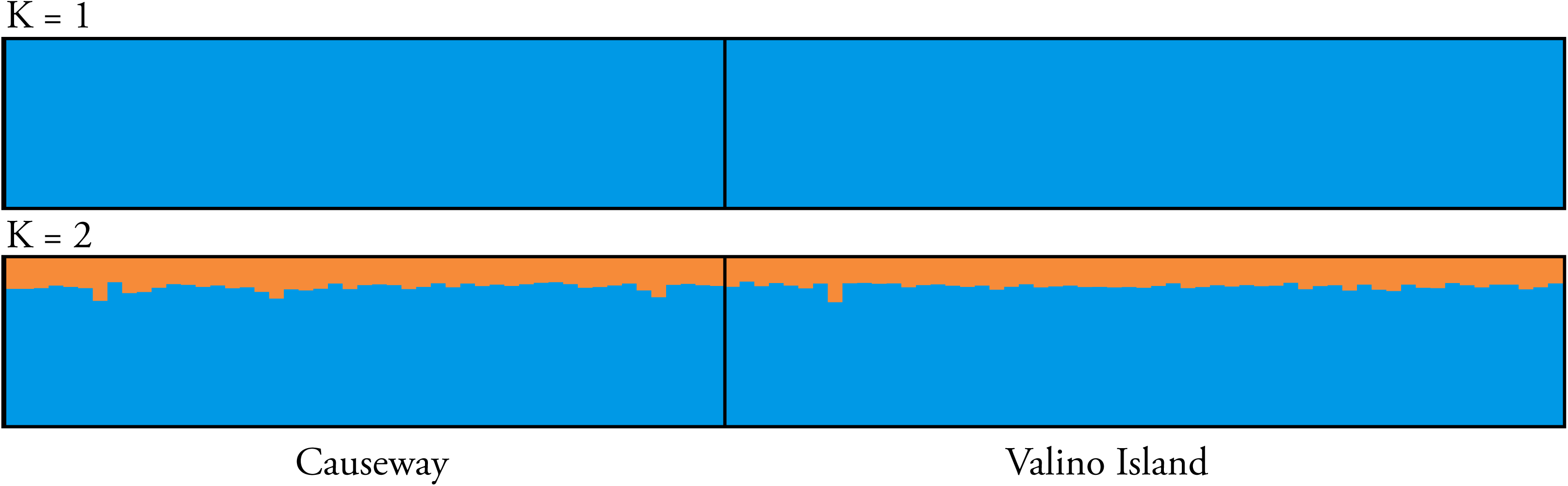
Results of STRUCTURE analysis, a Bayesian clustering method used to infer population structure. Each vertical bar represents an individual, with colors showing the proportion of that individual’s genome assigned to each genetic cluster for K = 1 and K = 2.

Genome-wide genetic divergence between the two collection sites was extremely low, with F_ST_ = 0.0074. In addition, there are no discernible regions of population separation along the genome (Fig. 04). To distinguish genuine population structure from sampling artifacts, we applied complementary permutation testing and bootstrap resampling approaches that examine different sources of variance in population genetic estimates.

**Figure 4.**
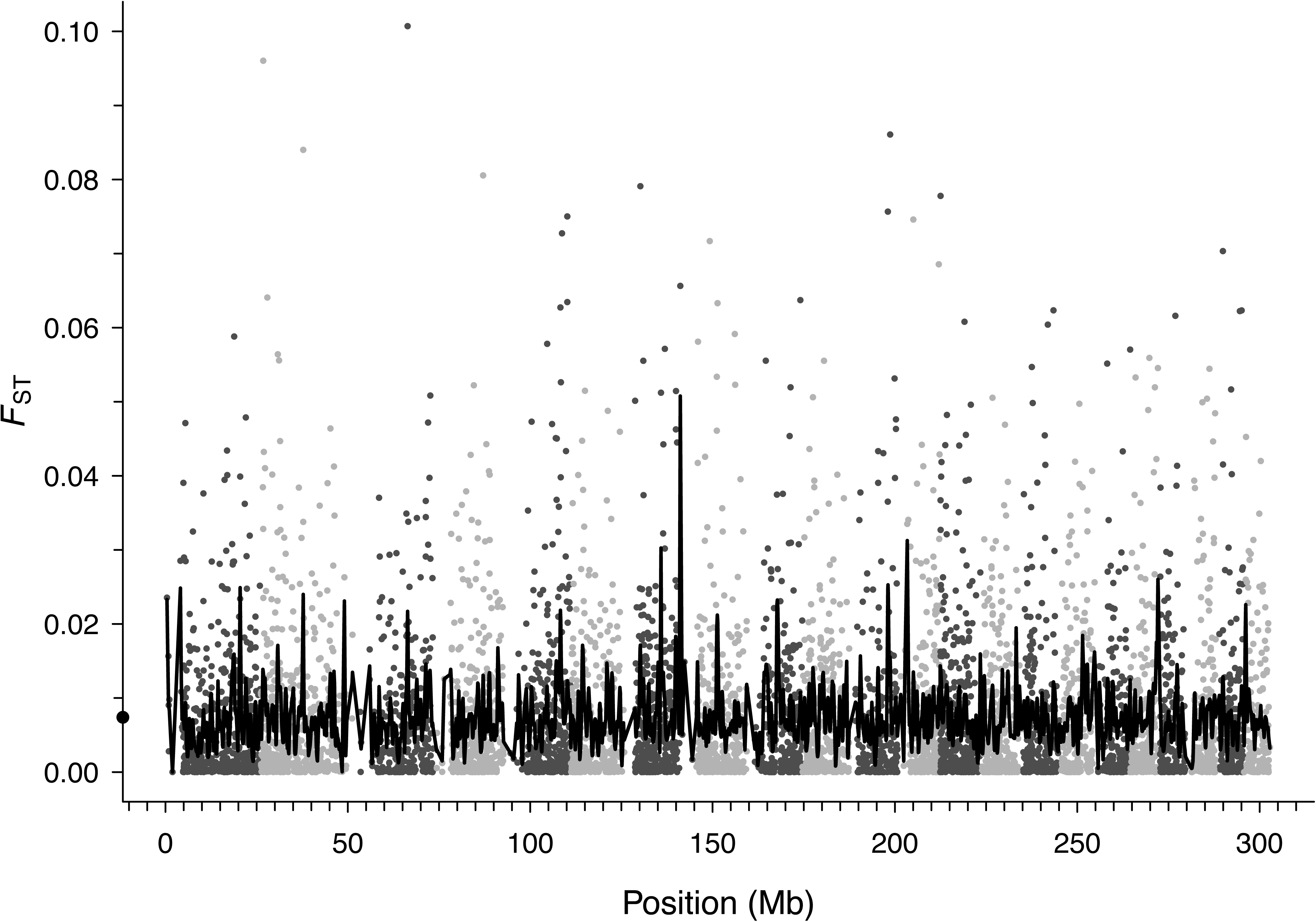
Genome-wide F_ST_ across chromosomes. Alternating shaded blocks represent successive chromosomes (1-22). Each point represents the F_ST_ estimate for an individual SNP, while the solid line indicates a smoothed trend across the genome. The black dot on the y-axis marks the genome-wide F_ST_ average of 0.0074.

Permutation testing evaluated whether the observed differentiation exceeded that expected under random population assignment. Across 100 permutation replicates, 99 permuted F_ST_ values met or exceeded the observed value, yielding a p-value of 0.990 and indicating that the observed differentiation could readily arise by chance (Supplemental Figure 2). Bootstrap resampling assessed whether differentiation could result from random sampling of loci. Using 1,000 replicates, each of which sampled 1,000 loci from the 6,074 SNP dataset, the bootstrap distribution had a mean F_ST_ of 0.0074 ± 0.0005 (95% CI: 0.0065–0.0083). The observed F_ST_ (0.0074) fell at the 49.3rd percentile, differing from the bootstrap mean by only 0.000008, demonstrating that observed differentiation lies well within expectations under random sampling of loci (Supplemental Figure 3). Together, these results indicate that the apparent genetic differentiation reflects sampling variance rather than systematic population structure.

These combined metrics provide robust evidence of panmixia between the two collecting locations. Further analysis treated the locations as one population.

### Large, Well-Mixed Population Exhibiting Low Kinship, Rapid LD Decay, and High Genetic Diversity

Kinship analysis indicated that most individuals were unrelated within and between sampling locations, with only occasional distant relationships (second-cousin level, kinship coefficient ≈ 0.027) (Supplemental Figure 4). At Valino Island, most coefficients were zero, with a few up to ∼0.028. Causeway showed a similar pattern, though slightly more frequent, reaching ∼0.036. Between locations, most pairs were unrelated, with a small subset exceeding the 0.027 threshold, indicating limited but detectable familial connectivity.

Genome-wide linkage disequilibrium (LD) analysis showed generally low levels of LD. Analysis of the relationship between LD metrics (both r² and D’) revealed a moderate overall correlation (r = 0.308, p < 0.001) across all intra-chromosomal SNP pairs within 5 Mb (Supplemental Figure 5 A). LD decayed rapidly outside of this interval, reaching background levels within 10.9 kb for r² and 7.0 kb for D’ (Supplemental Figure 5 B&C), indicating highly effective recombination across the *bay pipefish* genome.

A notable exception to this genome-wide pattern was a region on the distal end of chromosome 1 spanning the region from approximately 4 to 7 Mb (Fig. 5, A&B). This region exhibited substantially elevated linkage disequilibrium compared to genome-wide averages, with mean r² values ∼6 times higher (0.075 vs 0.012) and mean D’ values 1.78 times higher (0.550 vs 0.309) across 1,176 SNP pairs, consistent with reduced effective recombination in this 3 Mb segment. The elevated LD could reflect a misassembled region potentially due to repetitive sequence, a structural polymorphism, or could indicate the presence of a sex-determining region (McKinney et al., 2020). Further investigation is needed to distinguish between these possibilities.

**Figure 5.**
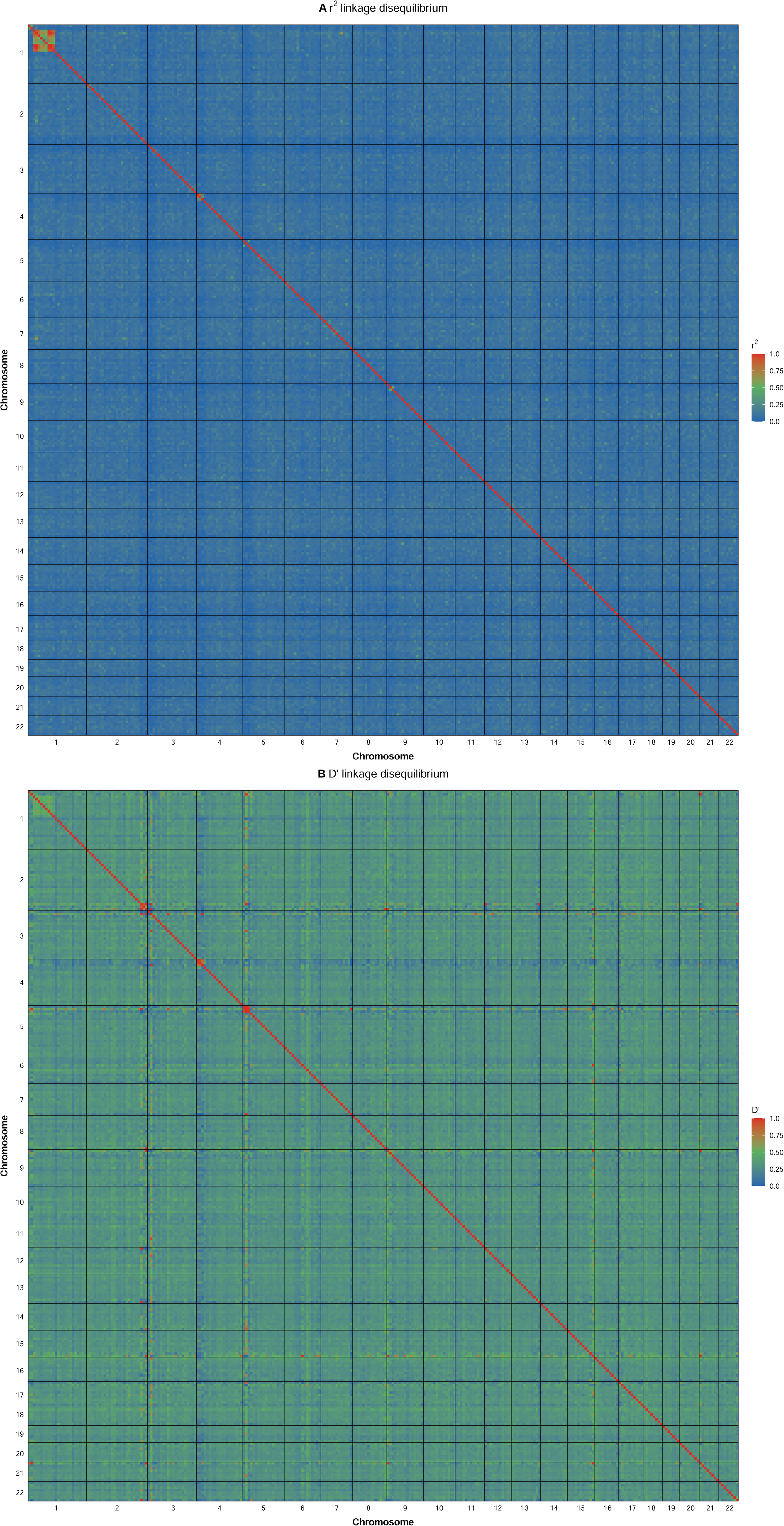
Genome-wide linkage disequilibrium (LD) patterns across 22 chromosomes. (A) Heatmap of maximum r² values and (B) heatmap of mean D’ values between 1 Mb genomic bins, highlighting intra- and inter-chromosomal LD. Black lines mark chromosome boundaries; the diagonal indicates perfect self-linkage (LD = 1). Color scales range from 0 (blue, no LD) to 1 (red, complete LD).

Estimates of contemporary effective population size (Ne) varied across two complementary methods. The allele frequency variance method produced the most precise estimate: Ne = 2,265 (95% CI: 2,207 - 2,329), based on 1,000 bootstrap replicates of 5,676 LD-pruned SNPs. The linkage disequilibrium method yielded a substantially higher estimate, Ne = 36,282 (95% CI: 11,230 - ∞), using 2,785 polymorphic markers and the recommended Pcrit threshold of 0.02.

These discrepancies reflect differences in methodological assumptions, and the temporal scales each approach captures. The variance method is sensitive to recent breeding dynamics, while the elevated LD-based estimate may reflect historical demographic events such as population expansion or ongoing gene flow from a larger metapopulation. Despite differences in magnitude, both approaches suggest that bay pipefish maintain an effective population size in the thousands, reinforcing the conclusion of a genetically diverse and panmictic population.

## DISCUSSION

While the number of high-quality reference genome assemblies available for syngnathids is growing (Ramesh et al., 2023; Scott-Somme et al., 2023; Wolf et al., 2024), population genomic assessments are lagging and are still needed to establish baselines for understanding connectivity and resilience in these vulnerable species. This is particularly true in estuarine systems like Coos Bay that are experiencing ongoing and increasing human influence. Given the limited population genomic information available for most syngnathid species, including bay pipefish, our study helps fill a key gap in understanding baseline patterns of genetic diversity and connectivity in this group.

In service of this goal, we produced the first genomic-scale assessment of genetic diversity, population structure, kinship, and effective population size in bay pipefish from Coos Bay, Oregon. Across multiple analyses, we observed patterns consistent with a large, well-mixed, and genetically cohesive population. Specifically, our results revealed high genetic diversity, minimal inbreeding, little to no population structure between sites, very low levels of relatedness among individuals, and low genome-wide linkage disequilibrium—together pointing to effective dispersal and gene flow within this estuarine system.

Population structure and kinship analyses collectively indicate that bay pipefish within Coos Bay are a single, well-mixed population. Individuals from the two sites showed no genetic clustering in multivariate analyses, and the very low genome-wide F_ST_ (0.0074) was statistically indistinguishable from random variation, suggesting that gene flow effectively prevents differentiation across the sites. Kinship analysis reinforced this conclusion, revealing that most individuals were unrelated, with no parent-offspring or full-sibling pairs detected and only occasional distant relationships observed. This pattern implies effective dispersal mechanisms or high population densities that limit the co-occurrence of close relatives. Large effective population size estimates further supported this interpretation: while the two methods of allele frequency variance and linkage disequilibrium differed in magnitude (Ne ∼2,265 vs. ∼36,000), both indicate that thousands of individuals contribute to reproduction, consistent with a demographically robust population capable of maintaining genetic connectivity across continuous estuarine habitat.

Bay pipefish have also shown variable genetic patterns across their range, depending on spatial scale and habitat context. Mitochondrial studies along the Pacific coast have suggested high gene flow among Oregon and California populations, though limited sample sizes constrained fine-scale resolution (Louie, 2003). A broader study identified significant genetic divergence between populations from Alaska, Washington, Oregon, and California, indicating that population structure emerges at larger spatial scales (Wilson, 2006). At finer scales, a study in Barkley Sound, British Columbia—where sites were separated by 9–83 km—found a mix of strong genetic divergence and panmixia among localities, shaped by habitat continuity and physical barriers like fjords (de Graaf, 2006). That study estimated a genetic neighborhood size of 40–60 km for *bay pipefish*, within which gene flow maintains genetic cohesion.

The two sites from which we sampled bay pipefish are separated by approximately 17 km of mostly continuous eelgrass habitat (Merkel & Associates, 2023), lacking significant physical barriers other than an industrial shipping channel, and our genetic estimates lie well within the genetic neighborhood size previously estimated for the species (de Graaf, 2006). These findings suggest that gene flow is ongoing at this spatial scale and that *bay pipefish* disperse effectively within continuous estuarine habitats, aligning with broader trends observed across the Syngnathidae and supporting the hypothesis proposed by Fritzsche (1980) that *bay pipefish* populations are genetically homogeneous within bays but may be differentiated across larger spatial scales. While our study is limited to two nearby locations within a single estuary, the patterns of panmixia, high diversity, and minimal kinship structure are consistent with this hypothesis. Future population genomic studies from other bays near Coos Bay can extend tests of Fritzsche’s predictions.

Previous population genetic studies in syngnathids have revealed a diversity of patterns, with some congeners of bay pipefish (the dusky pipefish*, S. floridae* and the gulf pipefish, *S. scovelli)*, maintaining high genetic diversity and fine-scale population structuring (Mobley et al., 2010; Flanagan et al., 2016) in a pattern of site fidelity and low dispersal, while the broadnosed pipefish, *S. typhle*, and the straight-nosed pipefish, *N. ophidion*, both associated with eelgrass habitats, show evidence of adult movement across eelgrass patches and weak site fidelity consistent with greater gene flow potential (Vincent et al., 1995). These contrasting patterns have been linked to differences in dispersal ability, larval duration, habitat fragmentation, and species-specific behavior–all of which can influence the spatial scale of gene flow and connectivity. In bay pipefish, panmixia might be aided by its large adult body size relative to many syngnathids, its large reproductive brood sizes, and the robust continuity of eelgrass habitat in estuaries like Coos Bay.

To more fully resolve the spatial genetic dynamics of bay pipefish, future studies should extend sampling to additional sites throughout Coos Bay and across other Pacific Coast estuaries. Broader spatial and temporal sampling, combined with focused studies of site fidelity and larval dispersal, will be essential for understanding the mechanisms that generate and maintain connectivity in this species. Our study helps establish a baseline for understanding genetic variation in bay pipefish in Coos Bay and provides a useful reference for future monitoring and comparisons across West Coast estuaries. More broadly, it contributes to our understanding of how spatial scale, habitat continuity, and life-history traits shape genetic structure in syngnathid fishes and offers guidance for future conservation and sampling efforts in estuarine systems.

## ACKNOWLEDGMENTS

Fish were collected under Oregon Department of Fish and Wildlife scientific taking permit number 26987. All animal research was conducted under University of Oregon Institutional Animal Care and Use Committee protocol AUP-20-23. Sequencing was performed at the Genomics and Cell Characterization Core Facility at the University of Oregon. This research was supported by the National Science Foundation grant IOS-2015301 to W.A.C. and by research funding from the University of Oregon.

## Author Contributions

M.C.C., M.A.W., V.G., and W.A.C. contributed to conceptualization and methodology. M.C.C., M.A.W., V.G., and W.A.C. conducted field investigations and specimen collection. M.C.C., M.A.W., V.G., and S.B. performed DNA extractions. M.C.C. constructed RAD-seq libraries, conducted formal analysis, created visualizations, and curated data. M.C.C. and M.A.W. wrote the original draft. All authors contributed to review and editing. W.A.C. acquired funding, provided resources, and supervised the project.

## DATA ACCESSIBILITY

Raw RAD-seq data generated in this study have been deposited in the NCBI Sequence Read Archive (SRA) under BioProject accession number PRJNA851781. The *Syngnathus leptorhynchus* reference genome has been submitted to NCBI GenBank (accession GCA_XXXXXXXXX, to be assigned upon release). Custom R scripts used for data analysis and figure generation are available on GitHub at https://github.com/wcresko/bay-pipefish-popgen.

