## Supplementary figures and images for "Genome-wide Analysis Reveals High Genomic Diversity and Panmixia in Bay Pipefish (*Syngnathus leptorhynchus*) from Coos Bay, Oregon"

### Supplemental Figure 1

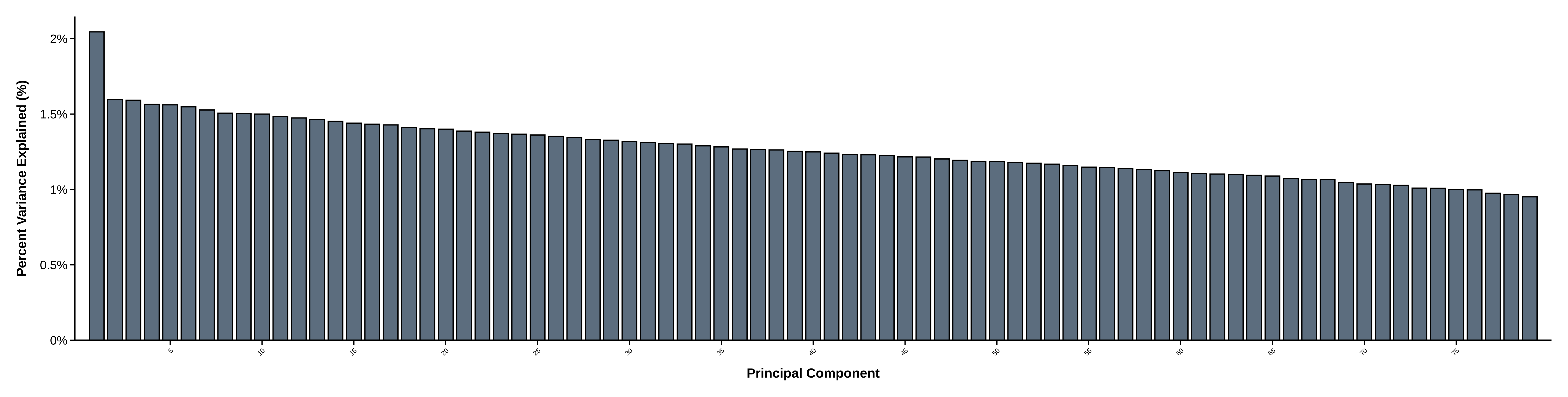

### Supplemental Figure 2

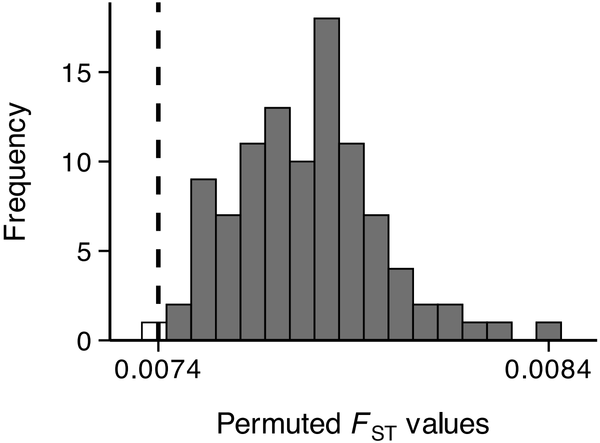

### Supplemental Figure 3

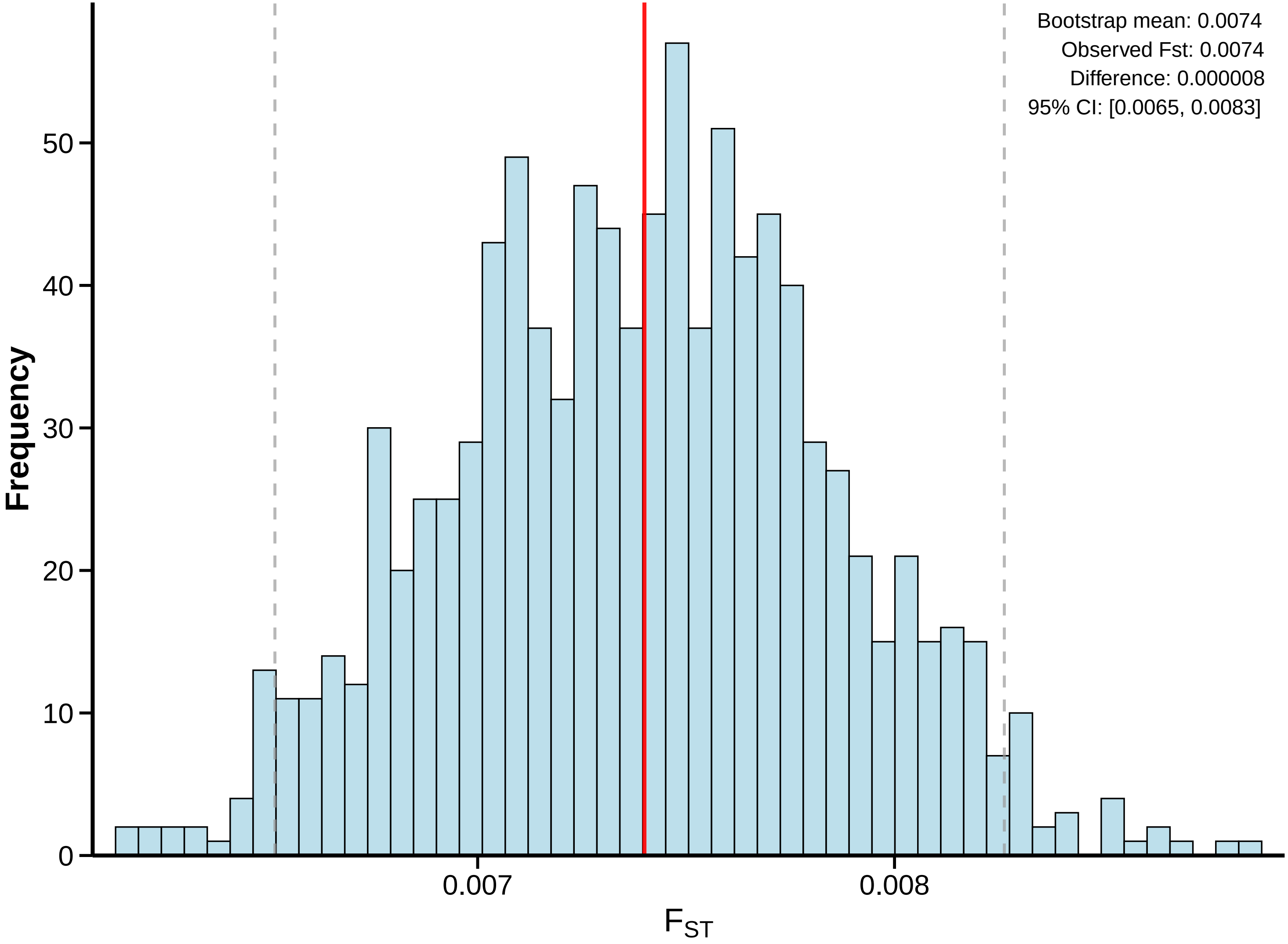

### Supplemental Figure 4

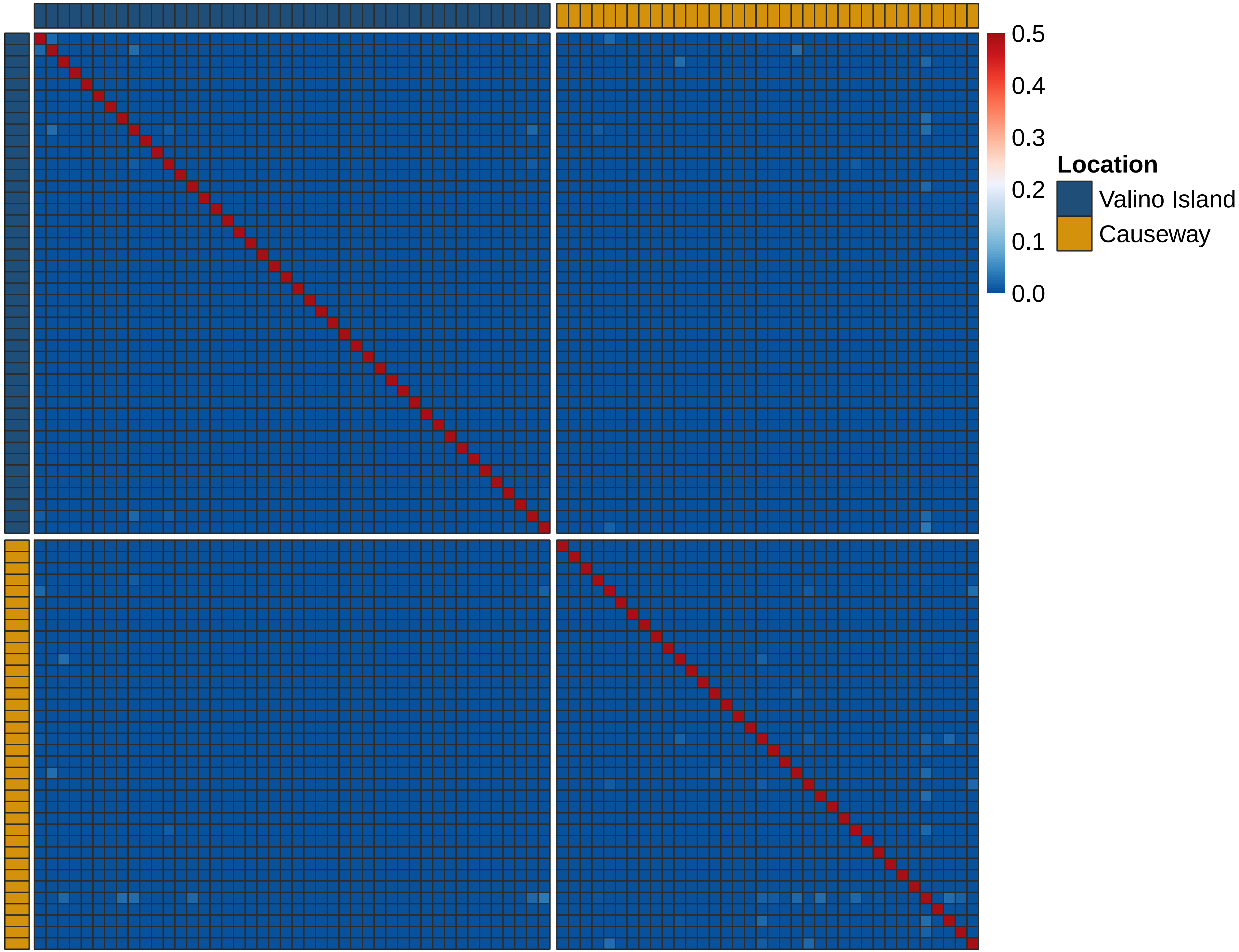

### Supplemental Figure 5

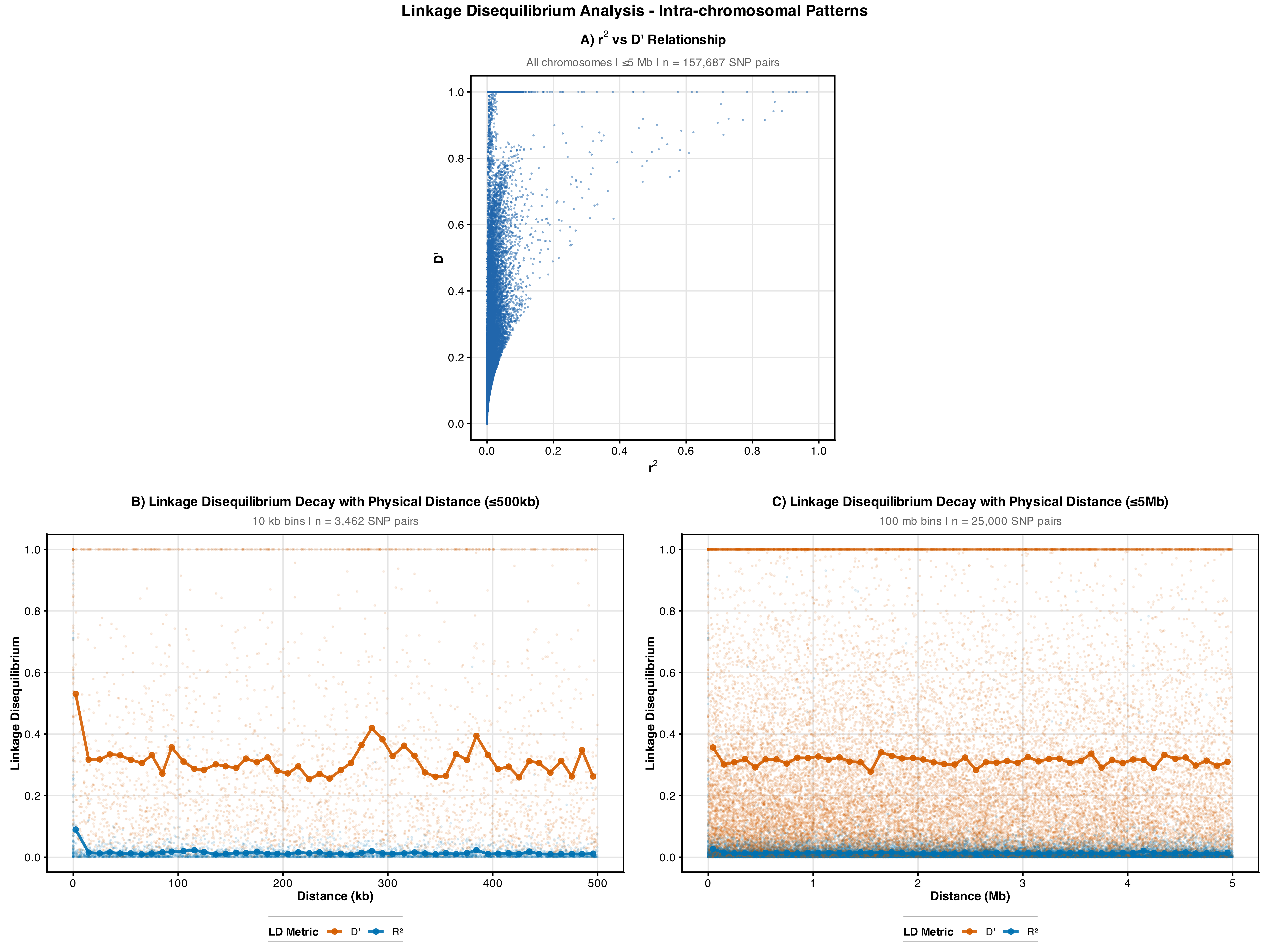
