## Supplemental Table 1 for "Genome-wide Analysis Reveals High Genomic Diversity and Panmixia in Bay Pipefish (*Syngnathus leptorhynchus*) from Coos Bay, Oregon"

**Supplementary Table 1. Summary statistics for the Illumina sequencing run after quality filtering.**

| **Illumina Run** | **Length** | **Total Sequences** | **Ambiguous Barcodes** | **Low Quality** | **Ambiguous RAD Site** | **Retained Reads** | **Percent Retained** |
| --- | --- | --- | --- | --- | --- | --- | --- |
| Slep | 135 | 501,953,205 | 102,205,346 | 76,030 | 26,527,08 | 373,144,745 | 74.3% |

This table presents sequencing quality metrics for the entire RAD-seq library (labeled "Slep") processed through the Stacks process_radtags module. The single row shows: Illumina Run name, read Length (in base pairs), Total Sequences obtained from the sequencer, and the number of reads removed due to Ambiguous Barcodes (no exact barcode match), Low Quality (Phred score <10 in sliding window), or Ambiguous RAD Site (no intact SbfI cut site). Retained Reads shows the total number of high-quality reads that passed all filters and were used for downstream analysis. Percent Retained indicates the overall proportion of raw reads that passed quality control. These metrics demonstrate the overall sequencing success and data quality for the complete dataset of 116 individuals.
