## Supplemental Table 2 for "Genome-wide Analysis Reveals High Genomic Diversity and Panmixia in Bay Pipefish (*Syngnathus leptorhynchus*) from Coos Bay, Oregon"

**Supplementary Table 2. Summary statistics generated by the** populations **module of Stacks for each sampling location.**

This table presents population genetic diversity measures for bay pipefish samples from two This table presents population genetic diversity measures for bay pipefish samples from two locations (Causeway and Valino Island) and combined. The table is divided into two sections: statistics calculated using only variant positions, and statistics calculated using both variant and fixed positions. **N** is the number of individuals retained after filtering. **Sites** refers to the total number of loci present in the dataset for each population. **Variant Sites** are the total number of segregating (polymorphic) SNPs detected. **Private** indicates the number of variant sites unique to that population across all loci. **H_EXP_** is the expected heterozygosity, **H_OBS_** is the observed heterozygosity, **Pi** represents nucleotide diversity (π), and **F_IS_** is the inbreeding coefficient (Weir and Cockerham's estimator); all are reported as means calculated across all loci. **Var** represents the locus‑wise (or site‑wise) variance of each statistic (H_EXP_, H_OBS_, π, F_IS_) across loci within the population.

| **Variant positions** | | | | | | | | | | | | |
| --- | --- | --- | --- | --- | --- | --- | --- | --- | --- | --- | --- | --- |
| **Pop ID** | **N** | **Sites** | **Variant**  **Sites** | **Private** | **H_EXP_** | **Var** | **H_OBS_** | **Var** | **Pi** | **Var** | **F_IS_** | **Var** |
| Causeway | 36 | NA | 6074 | 1 | 0.293 | 0.0196 | 0.257 | 0.021 | 0.298 | 0.020 | 0.130 | 0.059 |
| Valino Island | 44 | NA | 6074 | 9 | 0.297 | 0.019 | 0.259 | 0.019 | 0.301 | 0.019 | 0.136 | 0.053 |
| Combined | 80 | NA | 6310 | NA | 0.300 | 0.018 | 0.256 | 0.018 | 0.302 | 0.018 | 0.149 | 0.044 |
| **Variant and Fixed positions** | | | | | | | | | | | | |
| Causeway | 36 | 1486834 | 6074 | 1 | 0.001 | 0.0004 | 0.001 | 0.0003 | 0.001 | 0.0004 | 0.0005 | 0.0003 |
| Valino Island | 44 | 1486834 | 6074 | 9 | 0.001 | 0.0004 | 0.001 | 0.0003 | 0.001 | 0.0004 | 0.0005 | 0.0003 |
| Combined | 80 | 1513007 | 6310 | NA | 0.001 | 0.0004 | 0.001 | 0.0003 | 0.001 | 0.0005 | 0.000 | 0.0003 |
